# Multiplex immunohistochemistry of chronic active multiple sclerosis lesions links fibroblast-associated vessels with immune cell cuffs

**DOI:** 10.64898/2026.08.26.747283

**Authors:** R.P. Gorter, E. Liang, M. Goiko, V. W. Yong

## Abstract

**Background:** Multiple sclerosis (MS) is a chronic neurodegenerative disorder in which inflammatory demyelinating lesions affect the brain, optic nerve and spinal cord. MS lesion formation is accompanied by profound changes to blood vessels, including the density of PDGFRβ^+^ mural cells, historically identified as pericytes. Intriguingly, in recent years, single-cell and lineage tracing studies have shown that the PDGFRβ^+^ cell population is heterogeneous, comprising both pericytes and perivascular fibroblasts. Yet, due to their overlapping expression profiles, the spatial distribution of these cell populations in MS lesions remains poorly understood.

**Methods:** We employed multiplex immunohistochemistry for endothelial cells (CD31), basement membrane (laminin), fibroblasts (PDGFRβ, COL1A1, SMA), pericytes (PDGFRβ, SLC6A12) and immune cells (CD45, CD68) to characterize the spatial localization of fibroblasts and pericytes in MS lesions, and how this relates to perivascular space enlargement and immune cell presence.

**Results:** We analysed 17633 individual vessels across 5 control white matter, 5 normal-appearing white matter, 4 active and 4 chronic active MS lesions. By carefully delineating endothelium and perivascular compartments, we find that perivascular space area but not number of vessels is increased in MS lesions. Through mining of publicly available sequencing datasets, we confirm COL1A1 and SLC6A12 as fibroblast and pericyte markers, respectively, in the human brain. COL1A1^+^ and SLCA12^+^ vessels were largely distinct of one another. Unsupervised clustering of the expression profile of PDGFRβ, COL1A1 and SLC6A12 in individual vessels distinguished three partially overlapping vessel clusters. Of these, the fibroblast-associated vessel type (COL1A1 high, SLC6A12 low) was increased in chronic active lesion rim and center. Importantly, fibroblast-associated vessels were related to increased perivascular space enlargement and more accumulation of immune cells.

**Conclusion:** We identify distinct fibroblast- and pericyte-associated vascular phenotypes in human white matter. Notably, fibroblast-associated vessels are increased in chronic active lesions, where they are related to immune cell cuffs. These findings provide a spatial link between perivascular fibroblasts and chronic inflammation in MS.

## INTRODUCTION

Multiple sclerosis (MS) is a chronic neurodegenerative disease in which inflammatory demyelinating lesions affect the brain, spinal cord and optic nerve. The formation of these demyelinating lesions is accompanied by substantial changes to the vasculature^1^. On MRI, this can be appreciated by the central vein sign, i.e. the presence of a large vessel inside a white matter lesion^2^. At autopsy, MS lesions also show striking changes to the vasculature, including increased proliferation of endothelial cells^3^, elevated expression of basement membrane proteins such as laminins and collagens^4–6^ and changes in the density of PDGFRβ^+^ mural cells^7–9^. Historically, PDGFRβ^+^ perivascular cells in the brain have largely been considered pericytes, the cells in the brain that control capillary restriction and solute exchange^8^. However, in the era of single-cell resolution sequencing, there is increasing appreciation that the PDGFRβ^+^ cell population in the brain is heterogeneous and also comprises perivascular fibroblasts^10–13^.

Lineage-tracing studies combining PDGFRβ- and COL1A1-driven fluorescent reporters show that brain pericytes and fibroblasts are two distinct cell populations that differ with regards to morphology, marker expression and localization along the vascular tree^14^. For example, fibroblasts have a more flattened appearance, while pericyte cell bodies are rounder; fibroblasts express COL1A1, whereas pericytes do not, and fibroblasts are present in larger vessels, while pericytes mainly cover capillaries^14^. In line with this, single-cell/nucleus transcriptomics identifies distinct populations of pericytes and fibroblasts^15^, which have different expression profiles^16^. For example, fibroblasts are characterized by higher expression of ECM-related proteins, such as collagens and small leucine-rich proteoglycans^15^. Although fibroblasts in the healthy CNS are relatively rare, single cell/nucleus sequencing studies suggest that the fibroblast population expands following experimental demyelination^9–11,17,18^ and in human MS^18,19^. Yet, because of their overlapping expression profiles, examining the presence and spatial localization of fibroblasts versus pericytes within human MS tissue has thus far remained challenging.

Here, we used iterative antibody stripping and staining with MAXeraser^20^ to spatially characterize the expression of fibroblast and pericyte markers in control and MS white matter, and their association with immune cells. We identify fibroblast-associated, pericyte-associated, and mixed vascular phenotypes. We show that these vessels change in distribution across control white matter, normal-appearing white matter and MS lesions. Lastly, we identify a relationship between fibroblast-associated vessels and the perivascular accumulation of immune cells, relevant to understanding chronic inflammation in MS.

## METHODS

### Single-nuclei RNA-sequencing alignment and pre-processing

Raw data for the Absinta white matter dataset^19^ was retrieved from the SRA archive (PRJNA749443) and aligned using STARsolo as part of the nf-core scrnaseq pipeline (v2.1.0)^21^. STARsolo was run in the GeneFull mode to account for introns, with parameters set to the Cell Ranger 4 equivalent configuration, including the Cell Ranger version of EmptyDrops^22^. The data was pre-processed on a persample basis, with SoupX to correct for ambient RNA and scDblFinder to correct for doublets^23,24^. To remove outliers, low quality cells were first filtered using permissive mean absolute deviation (MAD)-based thresholds of 5 MADs for log1p of both total counts per barcode and number of genes per barcode and 3 MADs for the percent of mitochondrial counts per barcode. Cells with more than 5% mitochondrial counts or more than 5000 features were also removed. Retained features were required to be found in at least 2 cells. One chronic active lesion edge sample was removed from the analysis owing to low sample quality.

### Single-nuclei RNA-sequencing bioinformatic analysis

Pre-filtered sample count matrices were analyzed with Seurat v4 R package^25^. Normalization of counts was performed using SCT v2. on a per-sample basis. The samples were then rpca-integrated using 2500 variable features and a k.anchor of 5. The integrated assay was used to perform a dimensional reduction (principal component analysis, PCA) with 50 PCAs, followed by unsupervised clustering using FindNeighbors and FindClusters (resolution 0.5, algorithm 2). A UMAP was generated for the data using the integrated assay for cluster visualization. Differential expression analysis was performed using the Wilcoxon Rank Sum test on the recorrected SCT assay with FindAllMarkers and FindMarkers, with counts and data first re-corrected for multiple models using PrepSCTFindMarkers. Differentially expressed genes were identified on the basis of an adjusted p-value of < 0.05 following a Bonferroni correction to account for multiple comparisons and a Seurat v4 appropriate avg_log2(FC) cutoff. The cell type composition of clusters was identified based on canonical lineage markers and differentially expressed genes in each cluster.

### Human brain donors

Frozen brain tissue from 11 MS cases (average age = 52.7, range 39-82) and 5 non-demented controls (average age = 63.6, range 56-68) were obtained from the UK MS Tissue Bank at Imperial College, London (http://www.ukmstissuebank.imperial.ac.uk). All except two cases had progressive MS at time of death (Table 1). Participants had given informed consent for autopsy and use of materials for research.

**Table 1:**
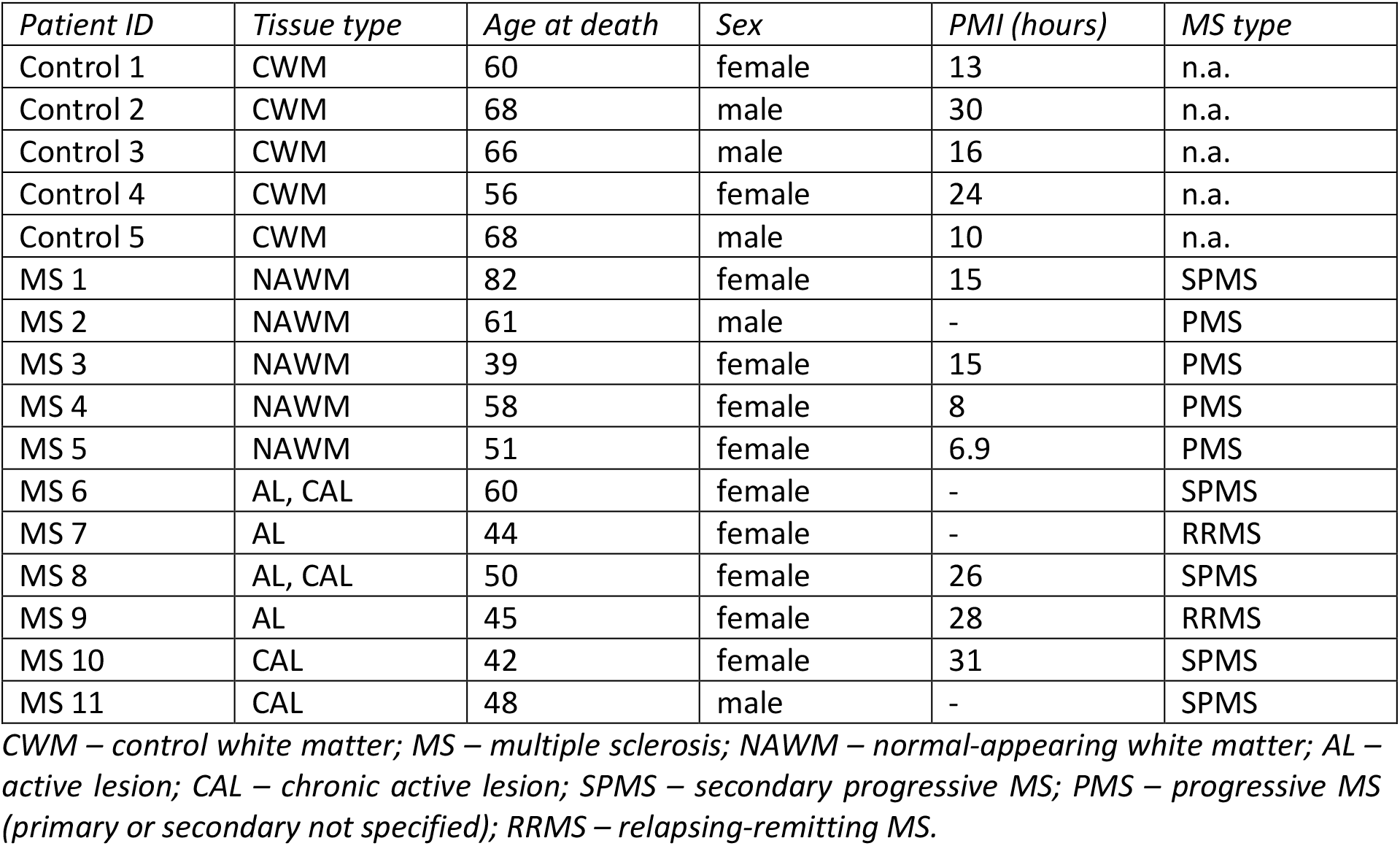
brain donor characteristics.

### Multiplex staining

To enable multiplex analysis of vascular and perivascular markers within the same tissue regions, 10µm frozen sections underwent four sequential rounds of immunofluorescence staining, imaging, antibody stripping, and restaining using MAXeraser^20^. The markers included in the present study were CD45 and SLC6A12 (round 1), CD68 and PDGFRβ (round 2), laminin 1+2 (round 3), and SMA, CD31 and COL1A1 (round 4). Antibody details and dilutions are provided in Table 1. For the first staining round, sections were air-dried for 15 min, fixed in 4% paraformaldehyde in PBS for 15 min at room temperature, and permeabilized with 0.25% Triton X-100 in PBS for 10 min. Sections were blocked for 1 h at room temperature in PBS containing 10% horse serum, 1% bovine serum albumin (BSA), 0.1% cold fish skin gelatin, 0.1% Triton X-100 and 0.05% Tween-20, followed by overnight incubation with primary antibodies at 4°C. The following day, sections were washed three times in PBS containing 0.1% Tween-20 and once in PBS. Tissue autofluorescence was quenched for 2 min using TrueBlack (Biotium, 23007) diluted 1:20 in 70% ethanol, followed by rinsing and washing in PBS. Sections were incubated for 2 h at room temperature with the corresponding fluorophore-conjugated secondary antibodies (1:500; Jackson ImmunoResearch Laboratories) and DAPI (1 µg/mL) in antibody dilution buffer. Sections were subsequently washed, mounted using Fluoromount-G (SouthernBiotech; 0100-01), and imaged using an Olympus VS120 slide scanner using the 20X objective. For subsequent staining rounds, coverslips were removed by incubating slides overnight in PBS in a custom 3D-printed horizontal slide holder, minimizing tissue damage during coverslip removal. Antibodies were stripped by incubating sections for 1 h at room temperature in MAXeraser solution, i.e. 30% m-Xylylenediamine (Sigma; X1202) and 3% sodium dodecyl sulfate (SDS; Sigma-Aldrich, L3771) in ddH_2_O, on a horizontal shaker at 50 rpm. Sections were washed three times in PBS, blocked for 1 h, and incubated overnight at 4°C with the next set of primary antibodies. Secondary antibody incubation, autofluorescence quenching, mounting, and imaging were performed as described above. The same tissue regions were imaged after each staining round. This stripping, restaining and imaging procedure was repeated for all four rounds.

### Confocal imaging

For simultaneous confocal analysis of CD31, PDGFRβ, SLC6A12 and COL1A1, 20 µm frozen sections were air-dried, fixed, permeabilized and blocked as described above. Sections were incubated overnight at 4°C with goat anti-PDGFRβ (R&D Systems, AF385, 1:50), together with primary antibodies against SLC6A12 and COL1A1 (Table 2). The following day, sections were washed and autofluorescence was quenched using TrueBlack as described above. Sections were then incubated for 2 h at room temperature with donkey anti-goat 594 (Jackson ImmunoResearch, 705-586-147, 1:500), donkey antirabbit 647 (Jackson ImmunoResearch, 711-606-152, 1:500) and donkey anti-mouse 790 (Invitrogen, A11371, 1:250). Following three washes in PBS, sections were incubated overnight at 4°C with directly conjugated sheep anti-CD31 (R&D Systems, AF806), which was attached to Alexa Fluor 488 using an Alexa Fluor® 488 Conjugation Kit Lightning-Link® (Abcam, ab236553). CD31 was directly conjugated to avoid cross-reactivity of the anti-goat secondary antibody with the sheep anti-CD31 primary antibody. The following day, sections were washed three times in PBS and mounted using Fluoromount-G. Images were acquired on a Leica STELLARIS 8 confocal microscope using a 10x objective.

**Table 2:** primary antibodies for multiplex analysis.

| *Antigen* | *Company* | *Catalogue number* | *Species* | *Concentration* |
| --- | --- | --- | --- | --- |
| SLC6A12 | Atlas Antibodies | HPA034973 | Rabbit | 1:1000 |
| CD45 | Invitrogen | MA5-17687 | Rat | 1:100 |
| PDGFRβ | Cell Signalling | 3169 | Rabbit | 1:50 |
| CD68 | Abcam | ab955 | Mouse | 1:500 |
| Laminin 1+2 | Abcam | AB7463 | Rabbit | 1:1000 |
| Smooth muscle actin | Abcam | ab124964 | Rabbit | 1:500 |
| CD31 | R&D Systems | AF806 | Sheep | 1:100 |
| COL1A1 | Sigma Aldrich | C2456 | Mouse | 1:500 |

### Image registration and analysis

Image analysis was performed in Qupath^26^, with registration of images using the Warpy extensions for QuPath and Fiji^27^. Images were manually aligned using DAPI staining, vascular landmarks, and other recognizable tissue features, followed by landmark-based registration and generation of a combined multichannel image in QuPath. A custom QuPath script was then used to identify CD31^+^ endothelium and laminin^+^ basement membranes and delineate the endothelial, perivascular, and, where present, intravascular compartments. Vessel segmentation was subsequently manually reviewed and corrected by an observer blinded to all channels other than CD31 and laminin. CD45^+^ and CD68^+^ cells were identified using trained QuPath cell classifiers, while positive area of PDGFRβ, SLC6A12, COL1A1 and SMA was determined using pixel thresholders. For each vessel and its corresponding compartments, vessel size, cell number, percentage of marker-positive pixels and mean fluorescence intensity were exported for subsequent analysis in R.

### Vessel analysis in R

QuPath object-level measurement tables were exported and processed in R to assign all vessel annotations to a tissue region and vessel unit. Analyses were performed using tidyverse packages for data handling and plotting, stats for k-means clustering and PCA, uwot for UMAP, cluster and mclust for clustering evaluation, and openxlsx for GraphPad-compatible Excel exports. Vessel density was calculated as the number of unique vessel units, i.e. a valid combination of CD31 and laminin, per mm^2^ tissue-region area. Total vessel area was calculated as summed vessel area (including intravascular space) relative to tissue-region area, and perivascular space enlargement as summed perivascular space area relative to summed endothelium + perivascular space area. Marker-positive pixel fractions were calculated as thresholded marker-positive area divided by the corresponding compartment area. For vessel clustering, only control white matter (CWM) and MS vessels with complete clustering features were retained: of 18298 vessel units, 17633 were included in the clustering analysis, corresponding to 3.63% excluded overall due to incomplete compartment/feature data. Regionspecific exclusions were 4.38% for CWM, 4.81% for normal appearing white matter (NAWM), 0.23% for active lesions, 0.67% for chronic active lesion rim, and 3.28% for chronic active lesion center. Of note, as a sensitivity analysis, clustering was also repeated including all vessels using median imputation for missing feature values, which produced highly similar cluster assignments and regional patterns. Clustering was performed in R using scaled vessel-level SLC6A12, PDGFRβ and COL1A1 meanintensity and positive-pixel-fraction features from endothelial and perivascular compartments. K-means clustering was performed with k=3, cluster labels were reordered by similarity of cluster centroids, and PCA was calculated from the same scaled feature matrix; UMAP was generated using all vessels for visualization. Cluster proportions were calculated as the percentage of vessels assigned to each cluster within each tissue region. Vessel-size distributions were summarized using total vessel area (including the intravascular space) and plotted on a log10 scale. For immune/perivascular space association analyses, CD45^+^ cell density was calculated within the endothelium + perivascular space compartment for each tissue-region-cluster and z-scored across clusters within the same tissue-region, so that 0 represents the unweighted tissue-region mean across clusters, with each cluster contributing equally regardless of vessel number. Perivascular space enlargement z-scores were calculated similarly using the mean ratio of perivascular space to endothelium area across vessels within each tissueregion-cluster. For this analysis, values were first calculated per vessel and then averaged within each tissue-region-cluster, preventing larger vessels from contributing disproportionately to the cluster-level estimate. Immune-cell distribution plots show the percentage of total tissue-region endothelium + perivascular space CD45^+^ cells assigned to each vessel cluster.

### Statistics in GraphPad Prism

Statistical analysis was performed in GraphPad Prism version 11.0.2. For comparisons between tissue regions, we used one-way-ANOVA with Dunnet’s post-hoc test, with CWM or NAWM as the reference group. When the assumption of equal variances was not met, we used Brown Forsythe and Welch ANOVA with Dunnet T3 test. For paired comparisons across three groups, such as CD45^+^ cell density across vessel clusters within the same samples, repeated-measures one-way ANOVA followed by Tukey’s multiple comparisons test was used, or Friedman test with Dunn’s post-hoc test for z-scored data. P < 0.05 was considered statistically significant.

## RESULTS

### Perivascular space area but not vessel number is increased in MS lesions

To characterize changes in perivascular fibroblasts and pericytes in MS, we first established the overall vascular architecture across control white matter (CWM), normal appearing white matter (NAWM), and MS lesions. Using a semi-automated workflow in QuPath, CD31 and laminin were used to delineate the endothelium, perivascular space (defined as the space between the endothelial and parenchymal basement membranes^28^), and, where present, the intravascular space (Figure 1A–F). In total, 18298 vessels were available for vessel-level analyses across five CWM samples (10157 vessels), five NAWM samples (3366 vessels), four active lesions (2201 vessels), and four chronic active lesions, in which rim (1203 vessels) and center (1371 vessels) were analyzed separately. As expected, the density of CD68^+^ cells was significantly increased in active lesions and chronic active lesion rims compared with CWM (Figure 1G). In contrast, vessel density varied considerably between lesions and did not consistently differ between CWM and MS tissue (Figure 1H). The percentage of tissue area occupied by vessels was only significantly increased in chronic active lesion centers (Figure 1I). However, the proportion of the vascular area occupied by perivascular space was consistently increased across lesion types, from an average of 23.9% in CWM to over 50% in active and chronic active lesions (Figure 1J). Together, these findings suggest that the apparent increase in vascular area in MS lesions is primarily associated with enlargement of the perivascular space rather than an increase in the number of vessels.

**Figure 1:**
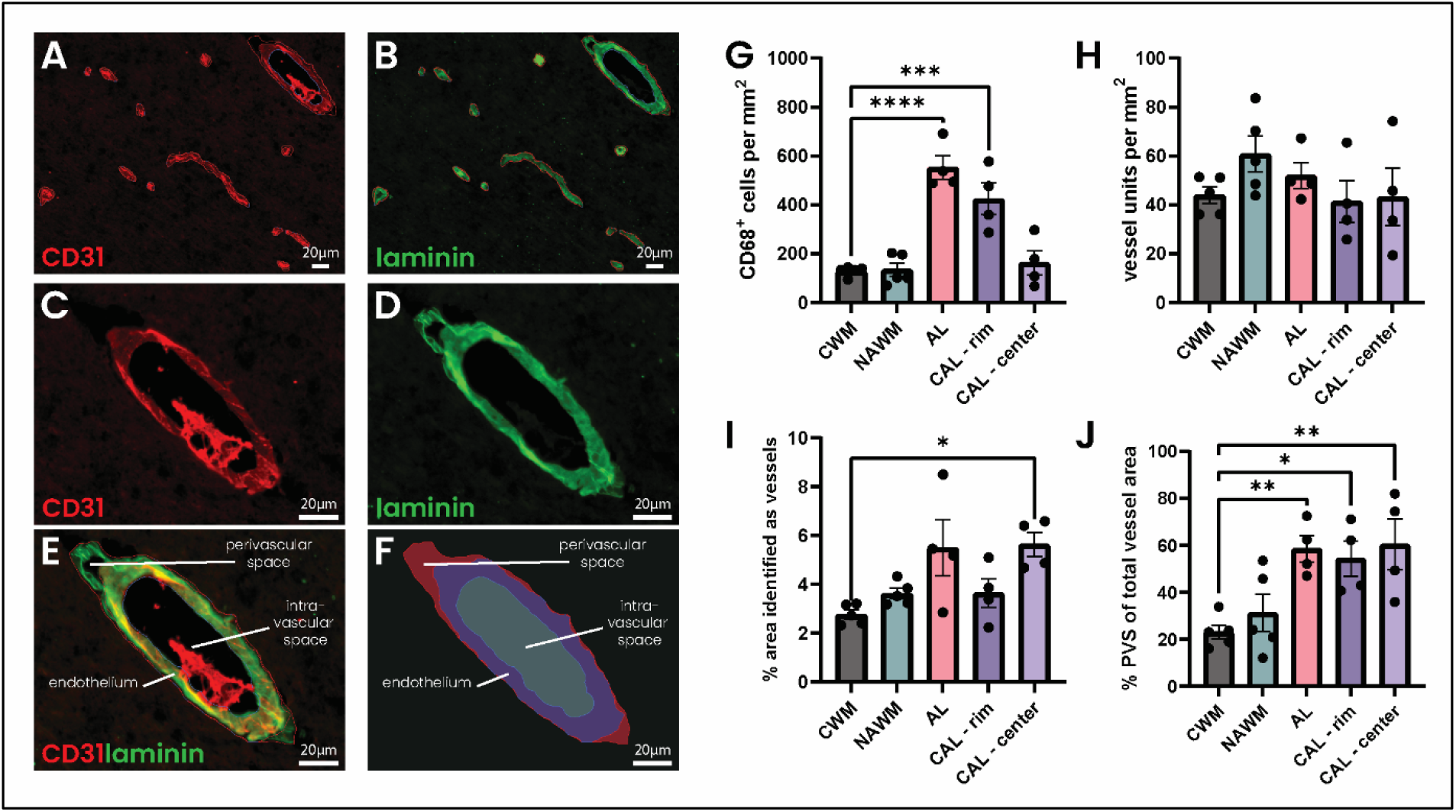
Perivascular space is increased in MS lesions without consistent changes in vessel density. Representative images of CWM showing (A) CD31 endothelial cell staining and (B) laminin for basement membranes with delineation of endothelial, perivascular and intravascular compartments in red. (C–E) Highermagnification images showing (C) CD31, (D) laminin, and (E) merged CD31/laminin staining, with arrows indicating the different vascular compartments. (F) Overlay showing the resulting compartment segmentation. (G) Quantification of CD68^+^ cells across regions (including CD68^+^ cells outside of vessels). (H) Density of vessel units, defined as valid combinations of CD31 and laminin, across regions. (I) Percentage of tissue area occupied by vessels (including the intravascular space). (J) Percentage of total vessel area occupied by perivascular space. Scale bars are 20 µm. AL – active lesion; CWM – control white matter; CAL – chronic active lesion; NAWM – normal-appearing white matter; PVS – perivascular space.

### SLC6A12 and COL1A1 distinguish pericytes and fibroblasts in the human brain

A recent study identified the solute carrier SLC6A12 as a specific marker of brain pericytes in CWM and Alzheimer’s disease^29^. To determine whether SLC6A12 also distinguishes pericytes in MS-affected white matter, we analyzed a publicly available single-nucleus RNA-sequencing dataset of CWM and chronic (in)active MS lesions^19^. Following quality control, 280 pericytes and 506 fibroblasts were identified. Across all cell types, SLC6A12 was differentially expressed in pericytes (log2FC > 0.25), whereas COL1A1 was differentially expressed in fibroblasts (Figure 2A–D).

**Figure 2:**
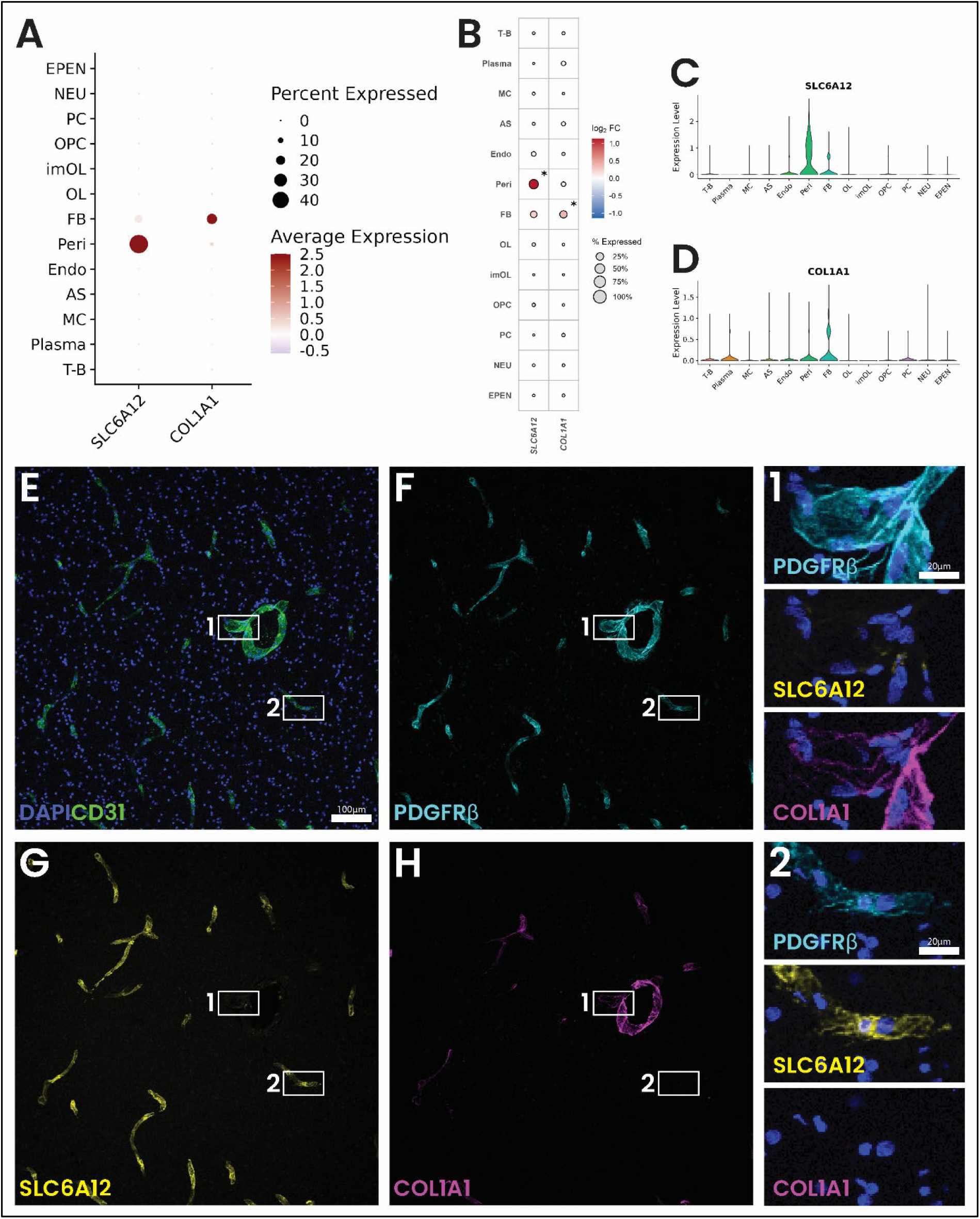
SLC6A12 and COL1A1 distinguish pericytes and perivascular fibroblasts in human white matter. (A)Dot plot showing expression of SLC6A12 and COL1A1 across cell types in a publicly available single-nucleus RNA-sequencing dataset of CWM and chronic (in)active MS lesions^19^. Dot size represents the percentage of cells expressing each gene and colour represents average expression. (B) Expression of SLC6A12 and COL1A1 within vascular and glial cell populations, with dot size representing the percentage of cells expressing each gene and colour representing log2 fold change. (C–D) Violin plots showing expression of (C) SLC6A12 and (D) COL1A1 across cell types. (E–H) Representative confocal images of CWM stained for (E) CD31, (F) PDGFRβ, (G) SLC6A12, and (H) COL1A1. (1–2) Higher-magnification images showing PDGFRβ, SLC6A12, and COL1A1 expression in representative (1) COL1A1^+^ and (2) SLC6A12^+^ vessels. CWM – control white matter; Epen – ependymal cells; FB – fibroblasts; Neu – neurons; OPC – oligodendrocyte precursor cells; imOL – immature oligodendrocytes; OL – oligodendrocytes; Peri – pericytes; Endo – endothelial cells; AS – astrocytes; MC – microglia; Plasma – plasma cells; T-B – T and B cells. Scale bars are 100um and 20µm, respectively.

We next assessed whether these markers distinguish the two populations at the protein level. Confocal imaging of CWM for CD31, PDGFRβ, SLC6A12, and COL1A1 showed PDGFRβ expression across CD31^+^ vessels, whereas COL1A1 and SLC6A12 showed largely mutually exclusive expression patterns. Vessels with high COL1A1 expression lacked SLC6A12, and vice versa. High-resolution imaging further revealed distinct cellular morphologies: COL1A1^+^ cells appeared flattened and elongated, whereas SLC6A12^+^ cells had a more rounded cell body (Figure 2). Together, these findings support the use of COL1A1 and SLC6A12 to distinguish perivascular fibroblast- and pericyte-associated vascular phenotypes, respectively, in MS white matter.

### Multiplex analysis identifies alterations in fibroblast and pericyte markers in NAWM and MS lesions

Next, to spatially characterize the distribution of perivascular fibroblasts and pericytes, we employed a stripping-restaining workflow^20^ to enable analysis of CD31, laminin, PDGFRβ, COL1A1, SLC6A12, SMA and CD45 in the same section across large tissue regions (Figure 3A–H). CD31 and laminin were used to delineate the endothelial and perivascular compartments (Figure 3A–D), after which expression of PDGFRβ, SLC6A12, COL1A1 and SMA was quantified within each compartment.

**Figure 3:**
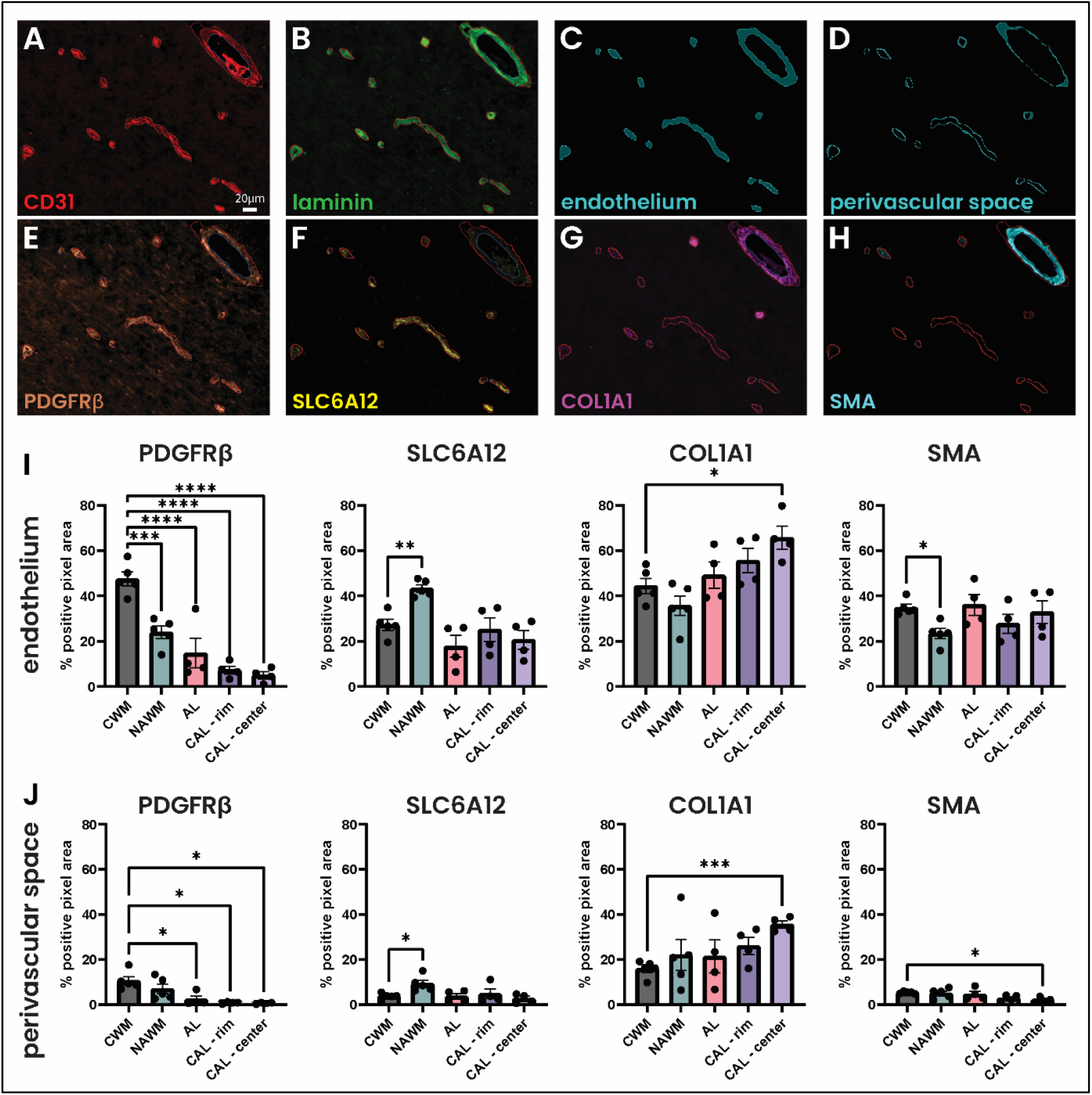
Pericyte- and fibroblast-associated vascular markers are altered in MS white matter and lesions. (A– H) Representative images showing multiplex staining and segmentation of vascular compartments. (A) CD31 and (B) laminin staining were used to delineate the (C) endothelium and (D) perivascular space. The same tissue region was analyzed for (E) PDGFRβ, (F) SLC6A12, (G) COL1A1 and (H) SMA. (I–J) Quantification of the percentage of positive pixels for PDGFRβ, SLC6A12, COL1A1 and SMA within the (I) endothelium and (J) perivascular compartments across CWM, NAWM, active lesions, and the rim and center of chronic active lesions. Each dot represents one tissue sample. Bars show mean ± SEM. *P < 0.05, **P < 0.01, ***P < 0.001, ****P < 0.0001. Scale bar is 20 µm. AL – active lesion; CAL – chronic active lesion; CWM – control white matter; NAWM – normalappearing white matter.

PDGFRβ, SLC6A12 and SMA expression was predominantly localized to the endothelial compartment, whereas COL1A1 also showed substantial expression within the perivascular space (Figure 3I–J). Compared with CWM, PDGFRβ expression was markedly decreased across NAWM and MS lesions, particularly within the endothelial compartment. In NAWM, endothelial SLC6A12 expression was increased, while SMA expression was decreased. In contrast, COL1A1 expression, a common marker for fibroblasts, increased toward chronic active lesions and was significantly elevated in the CAL center in both compartments. Together, these findings reveal substantial changes in pericyte- and fibroblastassociated markers across MS white matter, characterized by loss of PDGFRβ and increased COL1A1 expression with lesion progression.

### Unsupervised clustering of individual vessels identifies fibroblast-associated, pericyte-associated and mixed vessel populations

As changes within vascular subpopulations may be obscured when averaging marker expression across many vessels, we next examined pericyte- and fibroblast-associated markers at the level of individual vessels. After excluding vessels with incomplete data, expression profiles of SLC6A12, PDGFRβ, and COL1A1 within the endothelial and perivascular compartments of 17633 vessels were used for unsupervised clustering. This identified three partially overlapping vascular phenotypes, which were visualized by PCA and UMAP and showed distinct distributions across CWM, NAWM, and MS lesion regions (Figure 4A–E).

**Figure 4:**
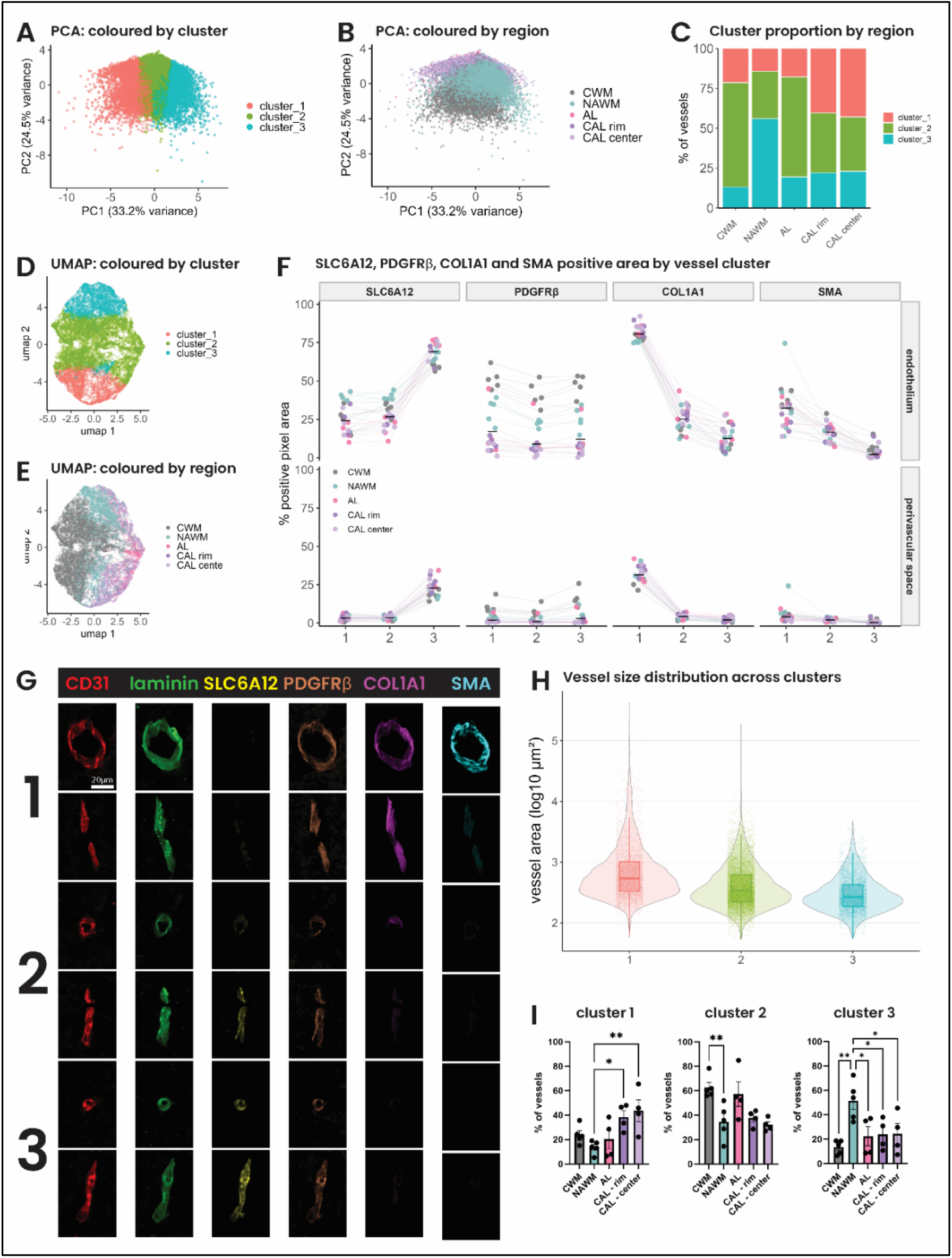
Unsupervised clustering identifies fibroblast-associated and pericyte-associated vascular phenotypes that are altered in MS. (A–B) PCA of individual vessels colored by (A) cluster and (B) tissue region. Percentages indicate the variance explained by principal components 1 and 2. (C) Proportion of vessels assigned to each cluster across tissue regions. (D–E) UMAP visualization of individual vessels colored by (D) cluster and (E) tissue region. (F) SLC6A12, PDGFRβ, COL1A1 and SMA expression in the endothelial and perivascular compartments according to vessel cluster and tissue region. SLC6A12, PDGFRβ and COL1A1 were included as clustering features, whereas SMA was analyzed independently. Lines connect mean expression values across clusters within the same tissue sample. (G) Representative images of vessels assigned to clusters 1–3 stained for CD31, laminin, SLC6A12, PDGFRβ, COL1A1 and SMA. (H) Distribution of vessel area across clusters. Violin plots show the distribution of individual vessels, with embedded boxplots indicating median and interquartile range. (I) Percentage of vessels assigned to clusters 1–3 within each tissue region. Each dot represents one tissue sample. Bars show mean ± SEM. *P < 0.05, **P < 0.01. AL – active lesion; CAL – chronic active lesion; CWM – control white matter; NAWM – normal-appearing white matter; PCA – principal component analysis. Scale bar is 20 µm.

Cluster 1 was characterized by high COL1A1 and low SLC6A12 expression, whereas cluster 3 showed high SLC6A12 and low COL1A1 expression (Figure 4F–G). Cluster 2 showed low to intermediate expression of both markers. PDGFRβ differed relatively little between clusters, although its expression was globally reduced in MS compared with CWM (Figure 4F). SMA was not included as an input feature for clustering but closely followed the pattern of COL1A1, with highest expression in cluster 1 (Figure 4F–G).

Vessel size varied considerably within all three clusters, consistent with the broad range of vessels analyzed, spanning from capillaries to large venules. However, cluster 1 contained larger vessels overall, with a median vessel area of 536 µm^2^ compared with 338 µm^2^ for cluster 2 and 265 µm^2^ for cluster 3 (Figure 4H). The relative abundance of the three phenotypes also differed across regions (Figure 4C,I). Cluster 1 was enriched in chronic active lesion rims and centers, comprising over 40% of vessels compared with only 21% in CWM. In contrast, cluster 3 was most abundant in NAWM, whereas cluster 2 was the dominant population in CWM and active lesions but appeared less frequent in NAWM and chronic active lesion regions.

Together, marker expression and vessel size support the identification of cluster 1 as a fibroblastassociated vascular phenotype and cluster 3 as a pericyte-associated phenotype, while cluster 2 represents a mixed phenotype. The increased abundance of fibroblast-associated vessels in chronic active lesions suggests increased involvement of perivascular fibroblasts in chronic MS pathology.

### Fibroblast-associated vessel cluster 1 is associated with immune cell accumulation and perivascular space enlargement

We next asked whether the identified vascular phenotypes differed in the extent of immune cell accumulation and perivascular space enlargement. We first quantified CD45+ cell density per vessel cluster across all tissue regions (Figure 5A). In addition, to account for differences in overall inflammation between tissue regions, CD45^+^ cell density within the endothelial and perivascular compartments was expressed as a z-score relative to the mean across clusters within the same tissueregion. Fibroblast-associated cluster 1 vessels showed the highest relative density of CD45^+^ cells, while mixed cluster 2 vessels were close to the regional average, and pericyte-associated cluster 3 vessels showed the lowest immune cell density (Figure 5A). Similarly, the perivascular space was relatively enlarged around cluster 1 vessels, but not around cluster 2 or 3 vessels (Figure 5B).

**Figure 5:**
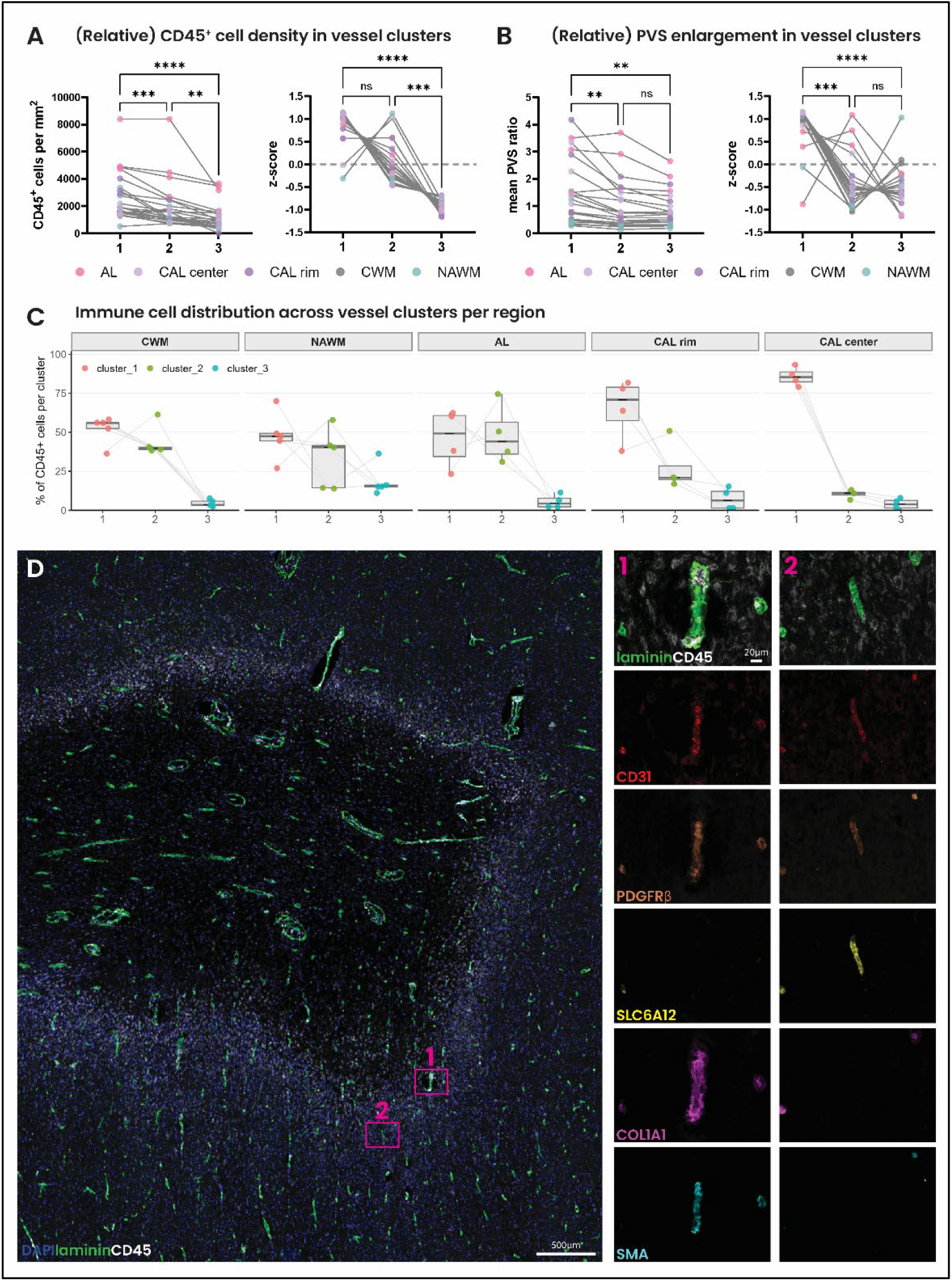
Fibroblast-like vessels are associated with immune cell accumulation and perivascular space enlargement. (A) CD45^+^ cell density across vessel clusters. Left, absolute density of CD45^+^ cells within the endothelial and perivascular compartments. Right, relative CD45^+^ cell density, expressed as a z-score across vessel clusters within each tissue region. A z-score of 0 represents the regional mean, with clusters weighted equally. A z-score of ±1 represents a value one standard deviation above or below the mean. Lines connect clusters from the same tissue sample. (B) PVS enlargement across vessel clusters. Left, mean ratio of PVS to endothelium area. Right, relative PVS ratio, expressed as a z-score across vessel clusters within each tissue region. A z-score of 0 represents the regional mean, with clusters weighted equally. A z-score of ±1 represents a value one standard deviation above or below the mean. Lines connect clusters from the same tissue sample. (C) Distribution of vessel-associated CD45+ cells across clusters 1–3 within CWM, NAWM, active lesions, and chronic active lesion rim and center. Values represent the percentage of vessel-associated CD45+ cells within each region assigned to vessels of each cluster. Lines connect clusters within the same tissue sample. (D) Representative image of a chronic active MS lesion showing laminin, CD45, and DAPI. (1–2) Higher-magnification images of a (1) cluster 1 vessel and (2) cluster 3 vessel stained for laminin/CD45, CD31, PDGFRβ, SLC6A12, COL1A1, and SMA. Scale bars, 500 µm (D) and 20 µm (1–2). AL – active lesion; CAL – chronic active lesion; CWM – control white matter; NAWM – normal-appearing white matter; PVS – perivascular space.

We next examined how vessel-associated immune cells were distributed across the three vessel phenotypes within each tissue region (Figure 5C). In CWM, approximately half of vessel-associated CD45^+^ cells localized to cluster 1 vessels. This association was more pronounced in chronic active lesions, where approximately 70% and 85–90% of vessel-associated CD45+ cells localized to cluster 1 vessels at the rim and center, respectively. Consistent with these quantitative findings, visual comparison of cluster 1 and cluster 3 vessels within the same lesion showed prominent perivascular immune cell accumulation around fibroblast-associated cluster 1 vessels, but not pericyte-associated cluster 3 vessels (Figure 5D). Together, these findings correlate the fibroblast-associated vascular phenotype with both enlargement of the perivascular space and accumulation of perivascular immune cells.

## DISCUSSION

In this study, we characterized changes in pericyte and fibroblast markers and their relationship to immune cells across control white matter and MS lesions. Through analysis of over 15000 individual vessels, we identified three main vascular phenotypes: a larger, COL1A1-high fibroblast-associated phenotype; a smaller, SLC6A12-high pericyte-associated phenotype; and a mixed phenotype. Intriguingly, the fibroblast-associated phenotype was increased in chronic active lesions, where it was correlated with enlargement of the perivascular space and accumulation of immune cells.

Previous immunohistochemical studies have already shown changes in the distribution of PDGFRβ^+^ cells in MS lesions^7–9^. However, PDGFRβ is expressed by both pericytes and fibroblasts, making it difficult to distinguish these populations using conventional immunohistochemistry. Here, analysis of multiple markers within individual vessels identified distinct fibroblast- and pericyte-associated expression patterns. Subsequent analysis showed that these phenotypes differ in vessel size, with fibroblast-associated vessels being substantially larger than pericyte-associated vessels. This corresponds well with previous work showing that fibroblasts are mainly found around arterioles and larger venules, while pericytes are predominantly associated with capillaries^14^. Together, the combination of marker expression and vessel size gives confidence that these clusters represent biologically distinct vascular phenotypes.

Fibroblast-associated vessels were particularly increased in chronic active lesion rims and centers, where they showed increased numbers of perivascular immune cells. Perivascular cuffing, defined by the accumulation of immune cells within the perivascular space, has long been recognized in MS pathology^30^. More recently, analysis across a large number of MS donors showed considerable heterogeneity in the extent of perivascular cuffing, with extensive cuffing associated with more severe disease^31^. Our finding that immune cells preferentially accumulate around fibroblast-associated vessels suggests that perivascular fibroblasts may be involved in the formation or maintenance of these cuffs. Of interest, immune cell-fibroblast interactions are well established in oncology^32^, but their relevance in the CNS is only beginning to be appreciated.

Our study has several limitations. Although a large number of vessels were analyzed, these originated from a relatively small number of donors, and future studies should validate these findings in larger cohorts. In addition, fibroblast- and pericyte-associated vessels were identified using a relatively small panel of markers. This is particularly relevant for the mixed phenotype, which could reflect an intermediate cellular state, but could also represent vessels containing both fibroblasts and pericytes. Additional markers or spatial sequencing approaches will be needed to better resolve these populations and determine the cellular interactions occurring within perivascular cuffs.

Overall, our findings show that perivascular fibroblast and pericyte phenotypes are altered in MS lesions, with fibroblast-associated vessels becoming more prominent in chronic active lesions and correlating with both perivascular space enlargement and immune cell accumulation. Better understanding how perivascular fibroblasts and immune cells interact within perivascular cuffs may provide insight into the mechanisms that sustain chronic inflammation in MS.

## ACKNOWLEDGEMENTS

We kindly thank Dr. Si-Hyung Park (Korea University College of Medicine) for their expert advice on MAXeraser stripping and Dr. Vincent Ebacher (Hotchkiss Brain Institute Advanced Microscopy Platform) for assistance with image alignment.

This study was funded by operating grants to VWY from MS Canada and the Canadian Institutes of Health Research. RPG acknowledges postdoctoral fellowships from MS Canada and the Dutch MS Research Foundation for the Gemmy and Mibeth Tichelaar Award.

